# The *expa1-1* mutant reveals a new biophysical lateral root organogenesis checkpoint

**DOI:** 10.1101/249961

**Authors:** Priya Ramakrishna, Graham A Rance, Lam Dai Vu, Evan Murphy, Kamal Swarup, Kamaljit Moirangthem, Bodil Jørgensen, Brigitte van de Cotte, Tatsuaki Goh, Zhefeng Lin, Ute Voβ, Tom Beeckman, Malcolm J Bennett, Kris Gevaert, Ive De Smet

**Author notes:** **CORRESPONDING AUTHOR**: Ive De Smet, VIB-UGent Center for Plant Systems Biology, Technologiepark 927, 9052 Gent, Belgium. Present address: Graduate School of Biological Sciences, Nara Institute of Science and Technology, Nara 630-0192, Japan (Tatsuaki Goh).

## Abstract

In plants, post-embryonic formation of new organs helps shape the adult organism. This requires the tight regulation of when and where a new organ is formed, and a coordination of the underlying cell divisions. To build a root system, new lateral roots are continuously developing, and this process requires asymmetric cell division in adjacent pericycle cells. Characterization of an *expansin a1* (*expa1*) mutant has revealed a novel checkpoint during lateral root formation. Specifically, a minimal pericycle width was found to be necessary and sufficient to trigger asymmetric pericycle cell divisions during auxin-driven lateral root formation. We conclude that a localized radial expansion of adjacent pericycle cells is required to position the asymmetric cell divisions and generate a core of small daughter cells, which is a prerequisite for lateral root organogenesis.

**SIGNFICANCE STATEMENT:** Organ formation is an essential process in plants and animals, driven by cell division and cell identity establishment. Root branching, where lateral roots form along the primary root axis, increases the root system and aids capture of water and nutrients. We have discovered that tight control of cell width is necessary to co-ordinate asymmetric cell divisions in cells that give rise to a new lateral root organ. While biomechanical processes have been shown to play a role in plant organogenesis, including lateral root formation, our data give new mechanistic insights into the cell size checkpoint during lateral root initiation.

## INTRODUCTION

In plants, organogenesis requires a well-defined sequence of (asymmetric) cell divisions (1). Especially during lateral root formation, the first rounds of pericycle cell divisions have been well characterized and are essential for the formation of a new lateral root organ (2–5). The first rounds of anticlinal pericycle cell divisions give rise to a lateral root initiation site (Stage I lateral root primordium) where small daughter cells are positioned next to each other in the center and are flanked by larger ones. The whole process of lateral root formation is regulated by several plant hormones, with auxin being the dominant signal (6–8).

During the formation of a new lateral root organ, a number of checkpoints have been defined, including auxin response oscillation in the basal meristem, migration of nuclei in adjacent pericycle cells toward the common cell wall, two rounds of consecutive and anticlinal asymmetric pericycle cell divisions followed by periclinal cell divisions of the small daughter cells, and auxin response in the endodermis (9–15). While these steps need to be correctly executed, at later stages, flexibility with respect to the number, order and orientation of cell divisions in the growing lateral root primordium is allowed (16, 17). Previously, it was shown that spatial accommodation by the endodermis is essential for the formation of a lateral root, and even for the first asymmetric cell divisions to occur (18). In addition, ablation experiments demonstrated that removal of the endodermis triggers unusual periclinal pericycle cell divisions, and auxin in the pericycle is required for correct anticlinal orientation of these divisions (19). Taken together, our current view of lateral root formation includes several checkpoints, but we still largely lack knowledge on how the different cells and cell types involved communicate. Furthermore, we do not know how all this information feeds into the necessary regulation of the cell wall, which is an active structure that plays a key role in cell expansion and is involved in several important physiological events.

Cell wall polysaccharides such as cellulose, hemicellulose and pectin form the major component of the primary cell wall in *Arabidopsis* (20, 21). Several models have been proposed for the architecture of the primary cell wall and its implications on wall extensibility (20, 22–25). The most recent ‘hotspot’ hypothesis proposes the presence of limited points of contact between the cellulose microfibrils mediated by xyloglucans that work as load-bearing sites and as targets of cell wall loosening (26, 27). There is also increasing evidence for the importance of pectin in control of wall extensibility, with formation of Ca^2+^-pectate cross-links considered to play a major load-bearing role in the absence of the cellulose-xyloglucan network in the cell wall (28–34). Although there is good understanding of the major components of the wall, the interactions between these components in an active cell wall and in response to developmental cues are not yet well understood. Alterations to the structure of the cell wall in response to growth are thought to be brought about by several cell wall remodeling agents that belong to different families and act upon different components of the cell wall (22, 35–38). Classic cell wall remodeling agents are expansins, which are known to alter the mechanical properties of the cell wall through non enzymatic, reversible disruption of non-covalent bonds in wall polymers, thereby creating local mechanical alterations in the wall (35, 39). Detailed characterization of the binding site of expansin in *Arabidopsis* cell walls highlighted the remarkable similarities in the expansin binding site to the ‘biomechanical hotspots’ described in the cell wall suggesting these sites to be the target sites of expansin action (26, 40, 27). However, the precise mechanism by which this occurs remains unclear.

Here, based on detailed analyses of the *expansin a1-1* (*expa1-1*) mutant, we define a novel lateral root organogenesis checkpoint. The *expa1-1* pericycle and lateral root initiation phenotypes revealed that a minimal pericycle width is necessary and sufficient to trigger asymmetric pericycle cell divisions during lateral root initiation.

## RESULTS AND DISCUSSION

### Meta-analysis identifies putative regulators of pericycle cell wall remodeling

During the early stages of auxin-triggered lateral root initiation, localized radial swelling of the lateral root founder pericycle cells takes place concomitant with the migration of the nuclei to the common wall (14, 18). Consequently, Stage I primordia showed a distinct bulging at the position of the small daughter cells (**Fig. 1*A-B***). Mechanistically, this likely involves (local) cell wall remodeling. To investigate the underlying molecular mechanisms associated with cell wall remodeling in the pericycle during lateral root initiation, we probed the differential expression of 406 putative cell wall remodeling agents in a previously published transcriptome dataset that was generated by sorting xylem pole pericycle cells that were synchronously induced by auxin for lateral root initiation and profiled for differentially expressed genes (15) (**Dataset S1**). This analysis identified 42 candidate genes that potentially play a role in pericycle cell wall modification accompanying the lateral root initiation process (**Dataset S1**). Given that plant hormones are key regulators of cell division and growth, also during lateral root formation, these candidate genes were evaluated against available transcriptome data sets to assess to what extent the expression of these putative cell wall remodeling agents is affected by plant hormones. We retained 15 candidates for which expression is regulated significantly by plant hormones (**Dataset S1**). These candidates were then compared against the list of potential key regulatory genes for the process of asymmetric cell division and cell fate specification during lateral root initiation (15) (**Dataset S1**). Based on this meta-analysis, we selected *EXPANSIN A1* (*EXPA1* - AT1G69530) for further characterization (**Fig. 1*C***).

**Fig. 1.**
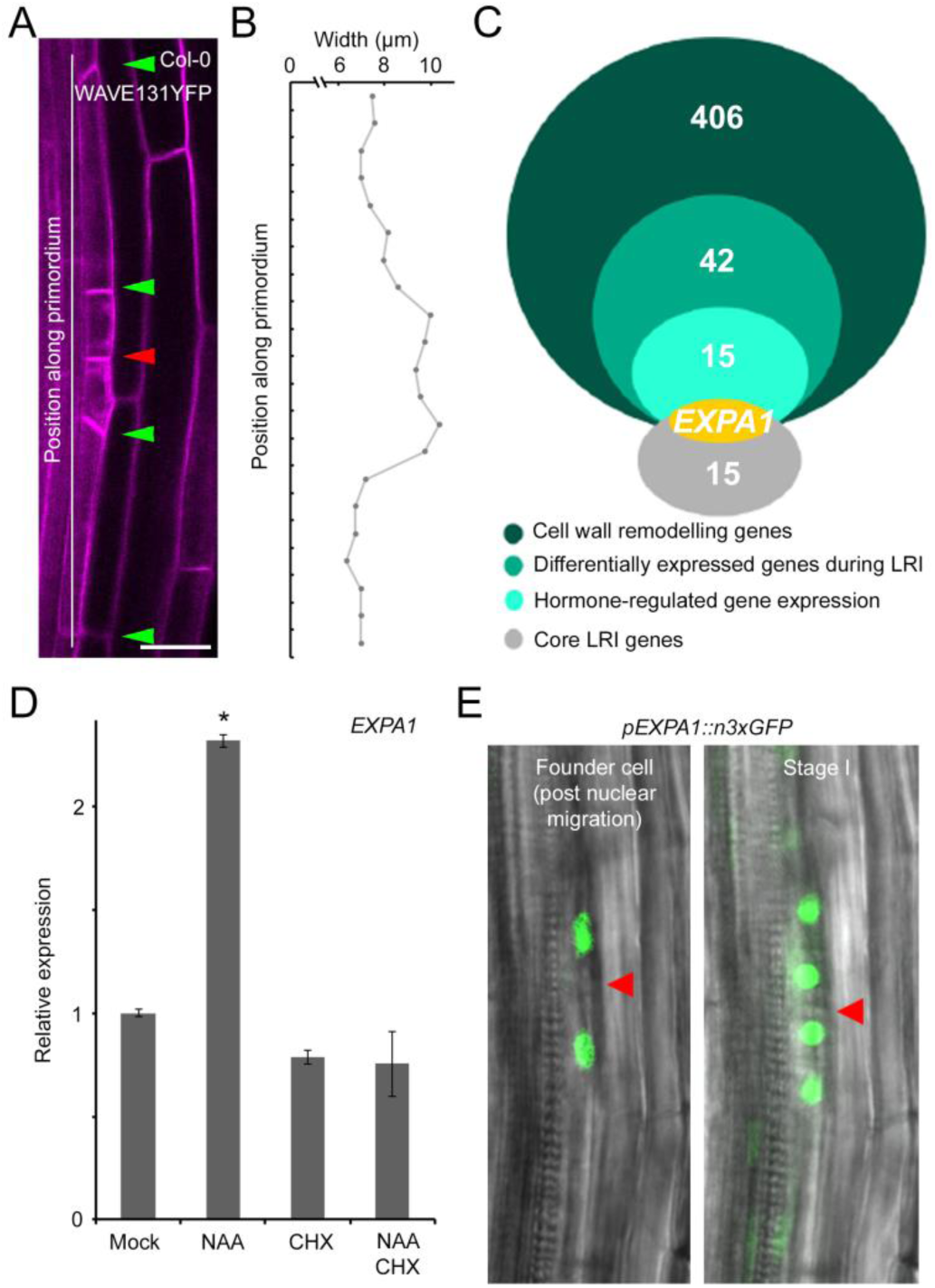
Meta-analysis identifies EXPANSIN A1 (EXPA1) as a potential regulator of lateral root initiation. **(A)** Representative image of Stage I lateral root primordium in Col-0.Cell outline visualized through the plasma membrane marker *pUBQ10::EYFP:NPSN12* (referred to as WAVE131YFP). Red and green arrowheads indicate the junction between adjacent pericycle cells that have divided asymmetrically and resulting daughter cells, respectively. Scale bar, 20 µm. **(B)** Representative pericycle cell width profile of Stage I lateral root primordium in Col-0. **(C)** Meta-analysis of genes encoding putative cell wall remodeling agents during lateral root initiation (LRI) leads to identification of *EXPANSIN A1* (*EXPA1*).**(D)***EXPA1* expression in Col-0 seedlings 3 days after germination transferred on mock, 10 µM NAA, 10 µM CHX or 10 µM NAA + 10 µM CHX plates for 6 h. Average of 3 biological replicates (with each ~15 roots) ± standard error. Statistical significance (Student’s t-test) compared with mock is indicated: *, p-value < 0.01. **(E)** Representative images of *invivo* expression analysis of *pEXPA1::n3xGFP* in the pericycle during lateral root initiation in lateral root founder cells post nuclei migration (left) and a Stage I lateral root primordium (right). Red arrowhead indicates junction between adjacent pericycle cells that undergo asymmetric cell division.

Since auxin has a dominant role in lateral root initiation (6), we initially focused on the regulation of *EXPA1* expression by this signal. We first confirmed that exogenous auxin treatment leads to an increase in *EXPA1* expression in the root (**Fig. 1*D*)**. To assess if this effect was direct, we used the protein synthesis inhibitor cycloheximide (CHX) in combination with auxin, as auxin-mediated degradation of labile AUX/IAA repressors results in ARF-regulated gene expression without *de novo* protein synthesis (41, 42). This revealed that *EXPA1* expression is regulated indirectly by auxin (**Fig. 1*D***).

To assess if *EXPA1* is specifically expressed in the root during lateral root initiation and to more directly connect EXPA1 activity with this process, we mapped the *EXPA1* expression pattern during the early stages of lateral root development. For this, we performed an *in vivo* expression analysis of *pEXPA1::n3xGFP* using a previously described bending assay (43). This revealed that *EXPA1* is expressed in the pericycle following nuclear migration preceding lateral root initiation, and then remained expressed in the small daughter cells (**Fig. 1*E* and Movie S1-2**). Apart from this, *EXPA1* expression was also observed in the cortical, but not in the endodermal cells overlying the lateral root primordium from Stage II onward (**SI Appendix**, **Fig. S1**). Taken together, the cell wall remodeling agent EXPA1 represents a novel component downstream of auxin-triggered lateral root initiation.

### Pericycle cell wall composition during auxin-induced lateral root initiation is affected in expa1-1

Since expansins play a role in cell wall loosening through altering interactions between the cell wall polymers (35, 40, 44), we investigated if loss of EXPA1 activity leads to changes in cell wall composition. For this, we identified an *expa1* T-DNA insertion line with very low *EXPA1* levels (about 5% of the control), named *expa1-1* (**SI Appendix, Fig. S2)**. First, we profiled key sugar monomers, namely glucose, galactose, arabinose and xylose, using High-Performance Anion Exchange Chromatography coupled with Pulsed Amperometric Detection (HPAEC-PAD) on hydrolyzed whole roots of Col-0 and *expa-1*. These sugar monomers are breakdown products of key cell wall polymers such as cellulose, xyloglucan and pectin polymers. HPAEC-PAD analysis revealed that *expa1-1* had reproducibly reduced galactose levels compared to Col-0 with no significant differences in other sugar monomers (**Fig. 2*A* and SI Appendix**, **Fig. S3-4**). Auxin has long been known to induce changes in the cell wall largely through acidification of the apoplastic space and triggering the activity of several cell wall remodelling agents, such as expansins (39, 45, 46). In order to evaluate if auxin-driven alterations in the cell wall require EXPA1, whole roots were treated with 10 µM 1-naphthalene acetic acid (NAA) for 6 h and analysed through HPAEC-PAD. Upon auxin treatment, no striking reproducible differences between wild type and *expa1-1* were observed; however, the galactose levels in *expa1-1* were restored to wild type levels (**Fig. 2*A* and SI Appendix**, **Fig. S4**). In parallel, we performed high-throughput Comprehensive Microarray Polymer Profiling (CoMPP) (47–49) on cell wall material of auxin-treated (6 h, 10 µM NAA) whole roots of Col-0 and *expa1-1*. However, we did not detect any major differences between the control and *expa1-1* using CoMPP (**Table S1**). Although the spatial resolution of the above-used techniques is poor due to whole root level analysis, we noted that the *expa1-1* roots exhibited altered cell wall polymers associated with galactose in their side chains, such as pectin and xyloglucan polymers (21, 48).

**Fig. 2.**
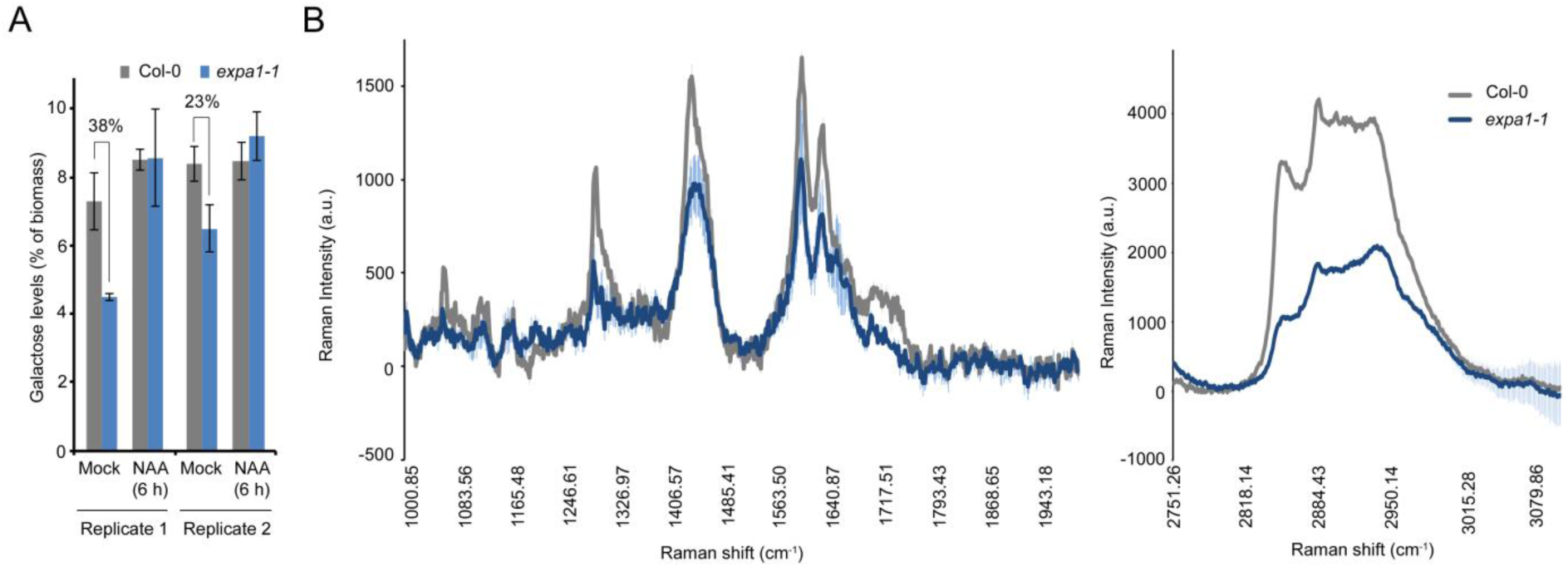
Cell wall composition of*expa1-1*. **(A)** Monosaccharide sugar analysis on hydrolyzed Col-0 and *expa1-1* roots at 7 days after germination with and without 6 h 10 µM NAA treatment using High Performance Anion Exchange Chromatography – Pulsed Amperometric Detection (HPAEC-PAD). Two independent biological replicates are shown, with average of technical replicates (2 or 3 for replicate 1 or 2, respectively) ± standard error. The % reduction is indicated. **(B)** Average of 4-6 measurements ± standard error depicting chemical spectra of pericycle cell junctions on cross sections of NAA-treated Col-0 and *expa1-1* roots in the range 1000 – 2000 cm^−1^ (left) and 2750 – 3100 cm^−1^ (right). a.u., arbitrary units. Statistical significance (Student’s t-test) compared is listed in **Table S2**: *, p-value < 0.05.

To investigate more subtle alterations in the biochemical composition of the pericycle cell wall at higher spatial resolution, we performed confocal Raman microscopy analyses on the xylem pole pericycle cell wall junctions during lateral root initiation. To probe the chemical structure of specific points of interest at high spatial resolution without the need to label or stain the sample, we used transverse sections of Col-0 and *expa1-1* of roots subjected to a previously described lateral root inducible system (15, 50–52) **(SI Appendix, Fig. S5-6**), and evaluated these for differences in their biochemical spectral profile. We observed changes in the biochemical spectral profile of the pericycle junctions in NPA-grown versus auxin-treated Col-0 seedlings (**SI Appendix, Fig. S7)**. In addition, we observed differences in the biochemical spectral profile of the pericycle junctions comparing NPA-grown Col-0 and *expa1-1* (**SI Appendix, Fig. S7**). But, most importantly, we observed that the intensities of several Raman bands for carbohydrate-based polymers, such as cellulose, xyloglucan and pectin, in auxin-treated Col-0 roots were less pronounced in auxin-treated *expa1-1* (**Fig. 2*B*)**. Overall, this would suggest that auxin-driven changes in the biochemical properties of the pericycle cell wall during lateral root initiation are altered in *expa1-1*, directly or indirectly. Nevertheless, our data suggest that EXPA1 is involved in pericycle cell wall remodelling upon auxin-induced lateral root initiation.

### EXPA1 controls patterned anticlinal asymmetric pericycle cell divisions

To evaluate if EXPA1 mediates lateral root initiation and/or emergence, we analyzed the *expa1-1* line. Taking into account that the primary root length is not dramatically different, a detailed lateral root staging analysis also did not reveal an obvious difference in any of the stages in the *expa1-1* mutant compared to the control (**Fig. 3*A* and SI Appendix, Fig. S8**). Therefore, to explore a possibly more subtle impact on lateral root initiation, we analyzed the progression through the stages of lateral root development using the bending assay (43). Examination of the root bends at 18 h post a gravitropic stimulus revealed a delay in the progression of the lateral root primordia from Stage I to Stage II in the *expa1-1* compared to the wild type (**Fig. 3*B*)**. To confirm that the observed lateral root initiation phenotype in *expa1-1* is due to the loss of EXPA1 activity, we performed complementation experiments. Indeed, expression of an *EXPA1* construct (*pEXPA1::EXPA1:6xHis*) in *expa1-1* could largely restore the lateral root Stage I to II progression defect in two independent lines (**Fig. 3*B***).

**Fig. 3.**
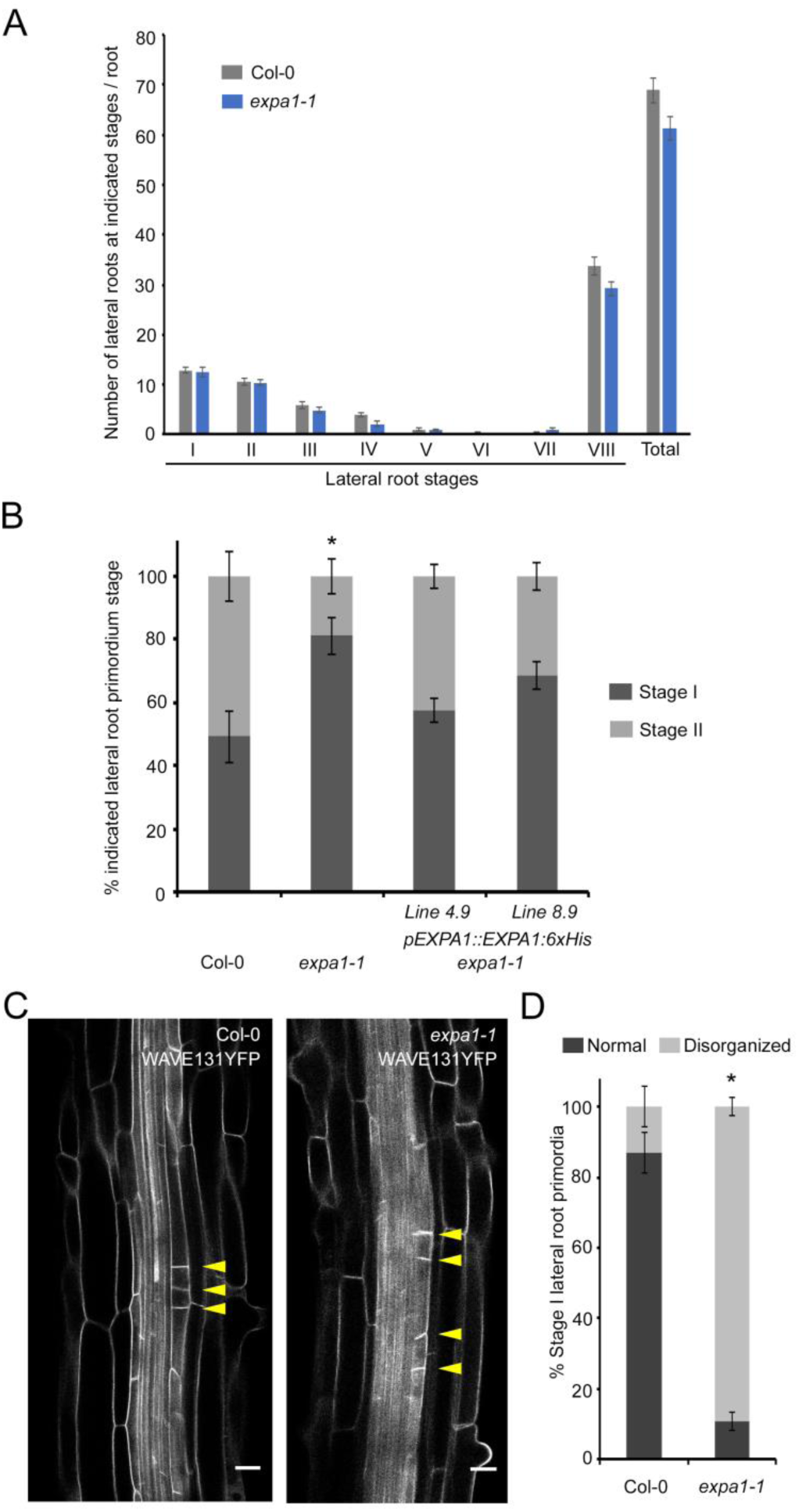
Lateral root initiation defects in *expa1-1*. **(A)** Lateral root staging of Col-0 and *expa1-1* roots at 10 days after germination for different stages of lateral root primordia. Average of 2 biological replicates (with ~6-8 roots each) ± standard error at p-value < 0.01. No statistical difference (using Student’s t-test) was found when compared with Col-0. **(B)** Progression through lateral root development using the bending assay at 18 h post bend in Col-0, *expa1-1* and two independent *expa1-1* lines expressing *pEXPA1::EXPA1-6xHis*. Average of combined results from 2-4 biological replicates each with 11-21 individual seedlings. Statistical significance (Student’s t-test) compared with Col-0 is indicated: *, p-value < 0.05. **(C)** Normal and disorganized Stage I primordia in Col-0 and *expa1-1*, respectively. Cell outline visualized through the plasma membrane marker *pUBQ10::EYFP:NPSN12* (referred to as WAVE131YFP). Scale bar, 20 µm.**(D)**Quantification of frequency of occurrence of normal and disorganized primordia in Col-0 and *expa1-1* at 10 days after germination. Average of 2 biological replicates ± standard error,with a total of 90 and 102 primordia from different Col-0 (7) and *expa1-1* roots (8), respectively. Statistical significance (Z Test Calculator for 2 Population Proportions, p < 0.05) compared with Col-0 (*) is indicated.

To explore if *expa1-1* displayed alterations in the pericycle cell division pattern during lateral root initiation, we introduced the plasma membrane marker line *pUBQ10::EYFP:NPSN12* (referred to as WAVE131YFP) (53) in *expa1-1* (referred to as *expa1-1* x WAVE131YFP) and performed a detailed *in vivo* analysis on the early stages of lateral root primordium development. In *expa1-1*, about 75% of what resembled Stage I primordia (which were included as Stage I in **Fig. 3*A***) showed small daughter cells not positioned next to each other, resulting in a loss of organized pattern (**Fig. 3*C-D*)**.

Given the lack of obvious differences in lateral root number and the early expression of *EXPA1,* we monitored the developmental progression of the early lateral root Stage 0 (pericycle cell on the root bend with no visible sign of division 12 hours post bend) to Stage II AI in *expa1-1* x WAVE131YFP. This revealed a transitionary lateral root development defect following the aberrant division pattern of Stage I primordia, as this appears to be compensated by subsequent divisions that restore a more or less organized Stage II lateral root primordium that continues to develop normally thereafter (**SI Appendix, Fig. S9**). Taken together, loss of EXPA1 activity affects the position of the first asymmetric and anticlinal cell divisions during lateral root initiation, resulting in the disruption of a typical Stage I pattern.

### EXPA1 is involved in pericycle cell expansion during lateral root initiation

Given the expected function of EXPA1, we analysed if there were any observable differences in the radial expansion of the xylem pole pericycle cells in *expa1-1*, which was measured as differences in cell width. This revealed a subtle difference between Col-0 and *expa1-1* (**Fig. 4*A-B***), but since only a limited number of pericycle cells are primed at any time to form a lateral root, pericycle cell width during lateral root initiation is not straightforward to capture and measure under regular growth conditions. Therefore, we subjected the roots of WAVE131YFP and *expa1-1* x WAVE131YFP to the above-mentioned lateral root inducible system that provides a uniform and synchronized population of pericycle cells undergoing lateral root initiation. Measurement of pericycle cell width of NPA-grown WAVE131YFP and *expa1-1* x WAVE131YFP roots, which a represent synchronously primed pericycle cell population that is not dividing, revealed that the pericycle cells of *expa1-1* x WAVE131YFP were significantly wider than WAVE131YFP (**Fig. 4*C-D* SI Appendix, Fig. S5)**. This is in agreement with the control conditions where we observed a subtle difference in pericycle cell width (**Fig. 4*A-B***). Upon auxin treatment, which induces lateral root initiation, the pericycle cells of WAVE131YFP increase in width (**Fig. 4*C-D***). Strikingly, the auxin-induced pericycle cell width of wild type pericycle cells corresponds to that of NPA-grown *expa1-1* pericycle cells (**Fig 4*C-D***), suggesting that *expa1-1* pericycle cells are at an advanced stage compared to wildtype. Furthermore, *expa1-1* pericycle cells did not increase further in width upon auxin treatment, and actually seem to reduce in width (**Fig. 4*C-D***), possibly explaining why there is no extensive division in these cells at this time point. In contrast, the width of endodermis and cortex cells seemed largely unaffected in *expa1-1*, compared to the control (**SI Appendix, Fig. S10**). In conclusion, loss of EXPA1 activity impacts pericycle cell width (**Fig. 4*A-D***), which appears to be essential for correct positioning of the anticlinal pericycle cell division (**Fig. 3*C-D***).

**Fig. 4.**
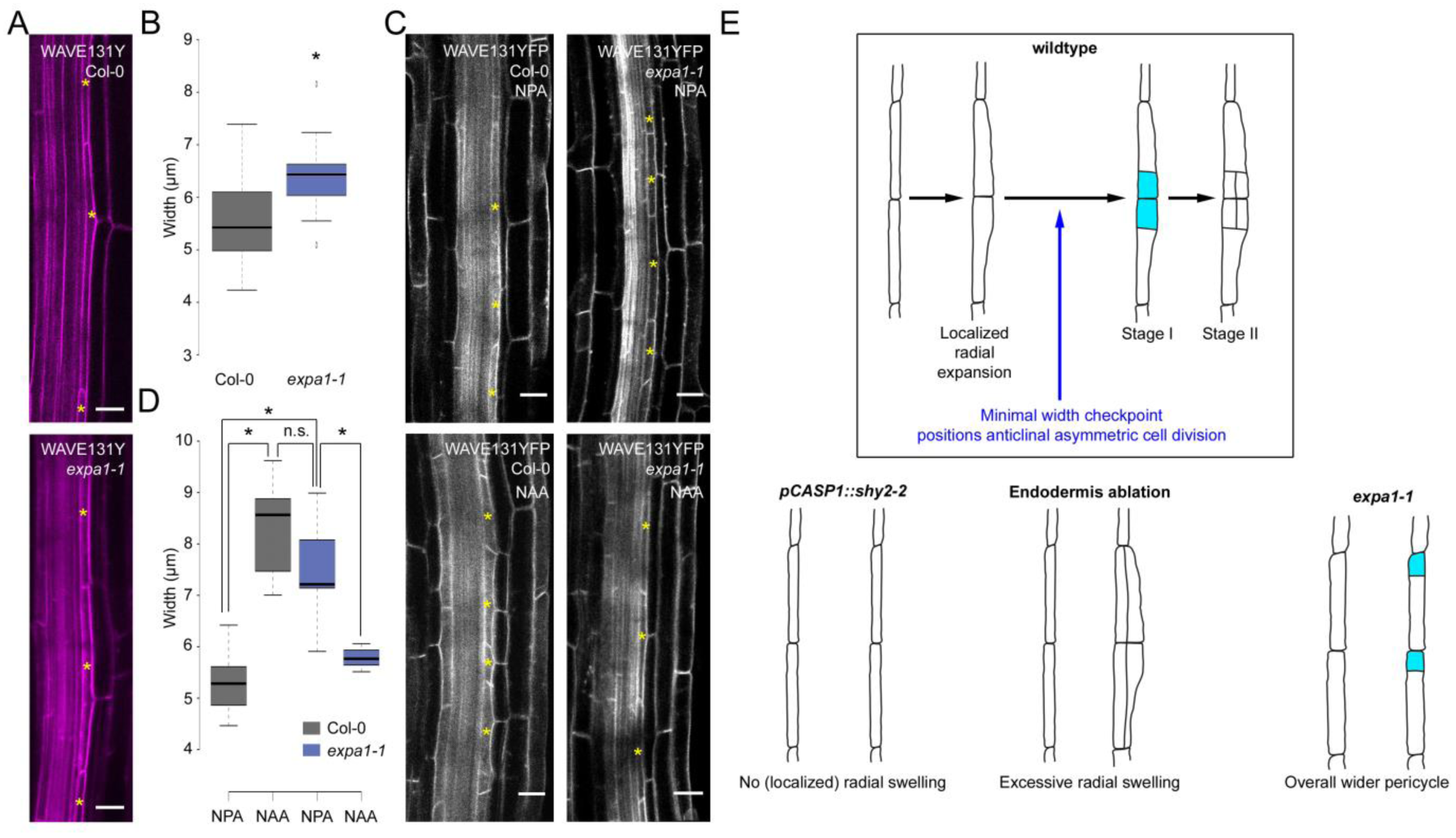
Pericycle cell width in *expa1-1*. **(A)** Representative images of Col-0 and *expa1-1* illustrating pericycle width in control growth conditions. Yellow asterisks mark pericycle cells. Cell outline visualized through the plasma membrane marker *pUBQ10::EYFP:NPSN12* (referred to as WAVE131YFP). Scale bar, 20 µm. **(B)** Boxplot of pericycle cell width in Col-0 and *expa1-1* in control growth conditions at 5 days after germination. At least 15 cells from 8 different roots were analyzed per genotype. Statistical significance (Student’s t-test) compared with Col-0 is indicated: *, p-value < 0.01. **(C)** Representative confocal images of Col-0 and *expa1-1* roots at 3 days after germination on 10 µM NPA or after transfer from NPA to 10 µM NAA for 6 h. Cell outline visualized through the plasma membrane marker *pUBQ10::EYFP:NPSN12* (referred to as WAVE131YFP). Yellow asterisk indicatesrepresentative pericycle cells used for cell width measurements. Scale bar, 20 μm. **(D)** Boxplot of pericycle cell width in Col-0 and *expa1-1* at 3 days after germination on 10 µM NPA or after transfer from NPA to 10 µM NAA for 6 h. At least 12 roots were analyzed per genotype and per treatment, with a total of at least 15 cells. Statistical significance (Student’s t-test) compared with Col-0 is indicated: *, p-value < 0.01.

## CONCLUSION

Several stages and checkpoints in early lateral root development have been described and auxin is central to this process (9–15). Here, through detailed characterization of the *EXPA1* expression pattern and *expa1-1* phenotypes in the pericycle, we could pinpoint a novel checkpoint in the early stages of lateral root development (**Fig. 4*E***). This biophysical checkpoint involving a minimum width requirement ensures that founder cells divide asymmetrically and anticlinally in an organized manner. If the pericycle width is too low as observed in the *pCASP1::shy2-2* line, no lateral root initiation takes place (**Fig. 4*E***) (18). If the pericycle width is too high, and no auxin-mediated correction occurs, periclinal cell divisions take place (**Fig. 4*E***) (19). In between, in wildtype conditions, pericycle cell swelling occurs on one side of two adjacent pericycle cells, resulting in the typical localized bulging of these cells and in organized anticlinal, asymmetric cell divisions (**Fig. 1*A* and 4*E***). If this widening takes place across the whole pericycle, as in the *expa1-1* mutant, the position of the anticlinal, asymmetric pericycle cell division does not take place in a coordinated way (**Fig. 4*E*)**.

The current understanding is that the expansins change the interaction between the wall polymer networks. EXPA1 is thought to target and alter ‘cellulose-xyloglucan’ junctions or hotspots in the wall, predominantly in dicots (26, 40), and this is important for cell wall loosening (31, 35). For example, suppression of *PhEXPA1* reduced the deposition of crystalline cellulose, resulting in thinner cell walls that were resistant to expansion (54). However, also binding to pectin has been observed, but this appeared unrelated to expansin activity (40). In the context of organogenesis, expansins were shown to alter the physical stress pattern in the meristem, so that tissue bulging occurs and cells achieve primordium identity (55, 56). At present, we cannot explain the counterintuitive phenotype in the *expa1-1* line with respect to cell wall composition and pericycle cell expansion. Whether the observed differences are a direct effect of the loss of EXPA1 activity, or a secondary effect resulting in an altered overall cell wall composition, remains to be investigated. Here, however, we mainly use the *expa1-1* mutant to define a novel checkpoint during lateral root organogenesis, and we are not assigning a direct role to EXPA1 in regulating pericycle cell wall composition.

In conclusion, altered EXPA1 activity impacts – directly or indirectly – on cell wall composition, which in turn controls pericycle cell width. During lateral root organogenesis, as is the case for expansins in other processes (57), the spatiotemporal pattern of *EXPA1* expression likely plays an important role and supports that EXPA1 also plays a direct role in the first stages of lateral root development. Finally, our analyses also revealed auxin-triggered changes in pericycle cell wall composition during early stages of lateral root development. In the future, it will be exciting to investigate what (polarity) cue drives local action of EXPA1, how this is linked to auxin in the pericycle cells and cell walls, how this is connected to nuclear migration in primed pericycle cells, and how this is coupled to the spatial accommodation by the overlaying endodermis.

## MATERIALS AND METHODS

Detailed materials and methods are described in **SI Appendix, Materials and Methods**.

### Plant materials and growth condition

*Arabidopsis thaliana* accession Columbia (Col-0) was used as wild type throughout the study. The T-DNA line for *EXPANSINA1* (AT1G69530), *expa1-1* (SALK_010506), was obtained from the Nottingham Arabidopsis Stock Centre (NASC)and genotyped using the following gene-specific primers: *CAAAGCAGACCACTATGACCC* and *TGTTCGGTAAGGCGTTGTTAG*. The *pUBQ10::EYFP:NPSN12* (WAVE131YFP) line has been previously described (53). All lines used for analysis were surface sterilized, stratified for two days at 4°C, and grown on vertical ½ MS plates [(2.154 g/l MS salts (Duchefa Biochemie MS basal salt mixture without vitamins), 0.1 g/l myo-inositol (Sigma), 0.5 g/l MES (Sigma), 10 g/l bacteriological agar (Sigma-Aldrich); pH 5.8 with 1M KOH)] at 22°C under continuous light conditions. For the 1-naphthaleneacetic acid (NAA) and cycloheximide (CHX) assay, 3 DPG (days post germination) roots were transferred onto vertical ½ MS plates supplemented with mock, 10 μM NAA, 10 μM CHX or 10 μM NAA + 10 μM CHX for 6 h and whole roots were analyzed. For the lateral root induction system-based analysis: WAVE131YFP and *expa1-1 x* WAVE131YFP lines were germinated and grown on 10 μM 1-N-naphthylphthalamic acid (NPA) for 72 h and transferred to ½ MS with 10 μM NAA for 6 h before analysis. Lateral root bending assays were performed on 3 DPG seedlings according to the method described previously (43).

### Microscopy

For localization/expression experiments, the Leica TCS-SP5 confocal microscope (Leica, Milton Keynes, UK) was used. Excitation and emission wavelengths were as follows: GFP – 488 and 485-545 nm; YFP – 514 and 525-600 nm; PI 514 and 570-670 nm. Roots were stained with 10 μg/ml propidium iodide (Sigma) for 2 min, rinsed and mounted in water.

### Confocal Raman microscopy

Col-0 and *expa1-1* seedlings were subjected to the lateral root induction system described above. See **SI Appendix, Materials and Methods** for details.

### Monosaccharide analysis

Whole roots of 7 DPG Col 0 and *expa1-1* seedlings grown on ½ MS at 22°C under continuous light condition were used for analysis. For auxin treatment, the 7 DPG seedlings were treated with 10 µM NAA in liquid ½ MS medium for 6 h followed by five washes in distilled water. The root fraction was isolated by cutting at the base of the hypocotyl and the samples were air dried at 37^°^C for two days. Dried root samples were hydrolyzed by Saeman hydrolysis (62) and monomeric sugar composition determined via High Performance Anion Exchange Chromatography with Pulsed Amperometric Detection (HPAEC-PAD). See **SI Appendix, Materials and Methods** for details.

### Comprehensive microarray polymer profiling (CoMPP) analysis

CoMP) analyses were performed as previously described (49). See **SI Appendix, Materials and Methods** for details.

## AUTHOR CONTRIBUTIONS

P.R., and I.D.S. designed research; P.R., E.M., L.D.V., K.S., G.A.R., K.M., B.v.d.C., T.G., B.J., K.S., and I.D.S. performed research; Z.L. contributed new reagents/analytic tools; P.R., U.V., T.B., M.J.B., and K.G. analyzed and interpreted data; and P.R. and I. D. S. wrote the paper.

## ACKNOWLEDGEMENTS

We thank Joop Vermeer for critical reading of the manuscript. P.R. acknowledges the financial support of the University of Nottingham Vice-Chancellor’s Scholarship for Research Excellence. The Confocal Raman microscopy work was supported by a University of Nottingham Interdisciplinary Centre for Analytical Science (UNICAS) award.

## SI SUPPLEMENTAL FIGURES

**Figure S1.**
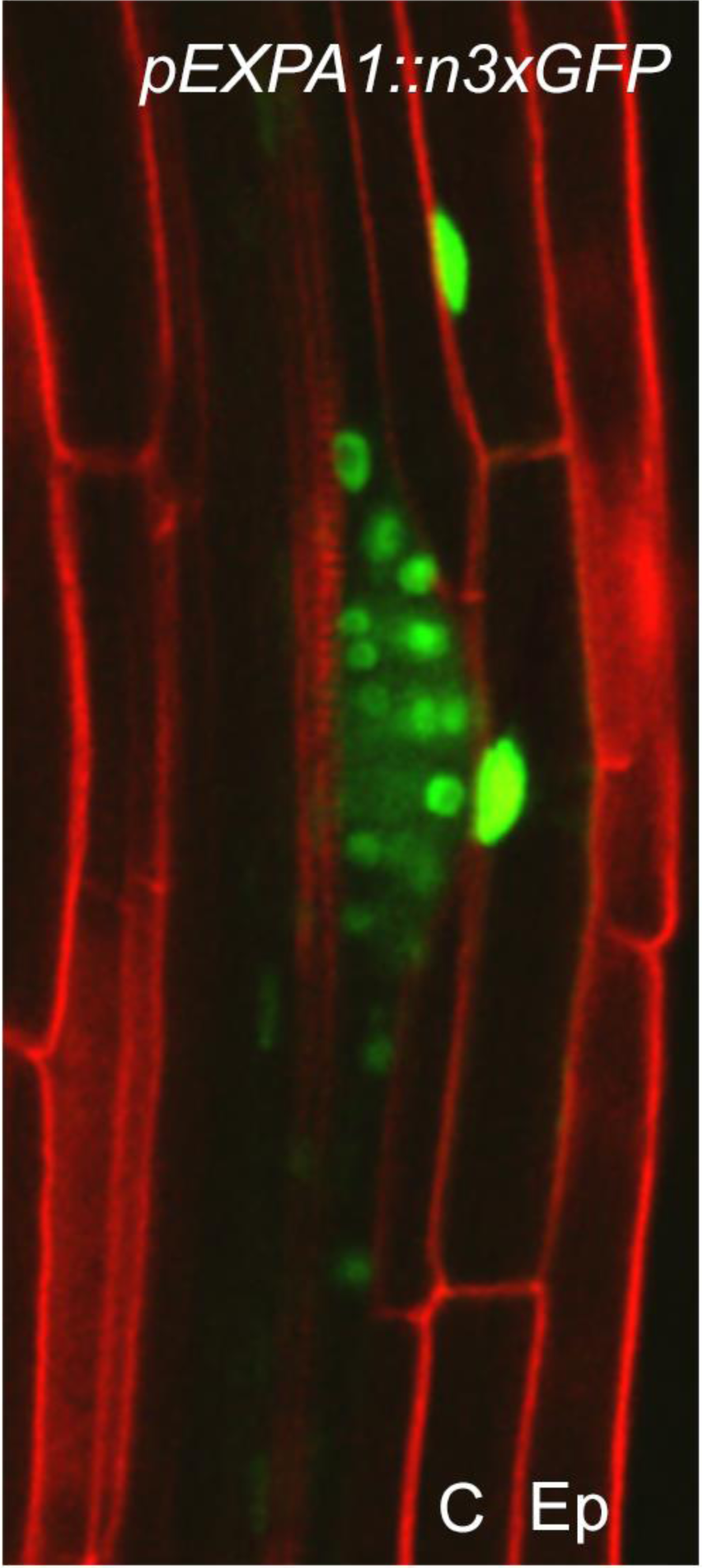
*pEXPA1::n3xGFP*expression in the cortical cells overlaying the lateral root primordium. C, cortex; Ep, epidermis. Cell outline visualized by staining with propidiumiodide.

**Figure S2.**
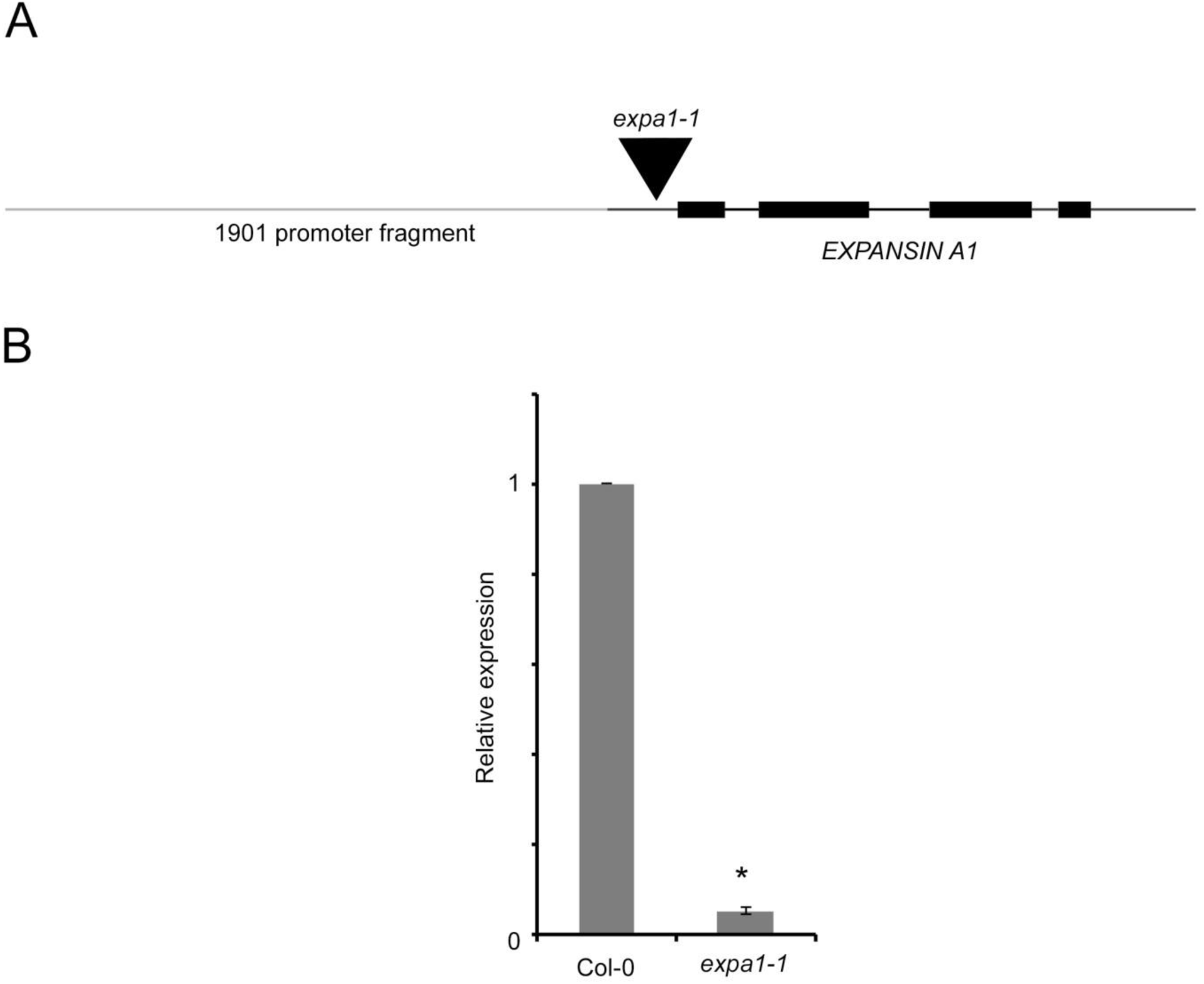
(**A**) Schematic representation of the*EXPA1*genomic region, indicating the position of the T-DNA insert (60 bp upstream of the transcriptional start site in the 5′ UTR region of *EXPA1*). UTRs, dark grey lines. **(B)** qRT-PCR analysis of *EXPA1* expression in *expa1-1*. Average of 3-4 biological replicates ± standard error. Statistical significance(Student’s t-test) compared with Col-0 is indicated: *, p-value < 0.01.

**Figure S3.**
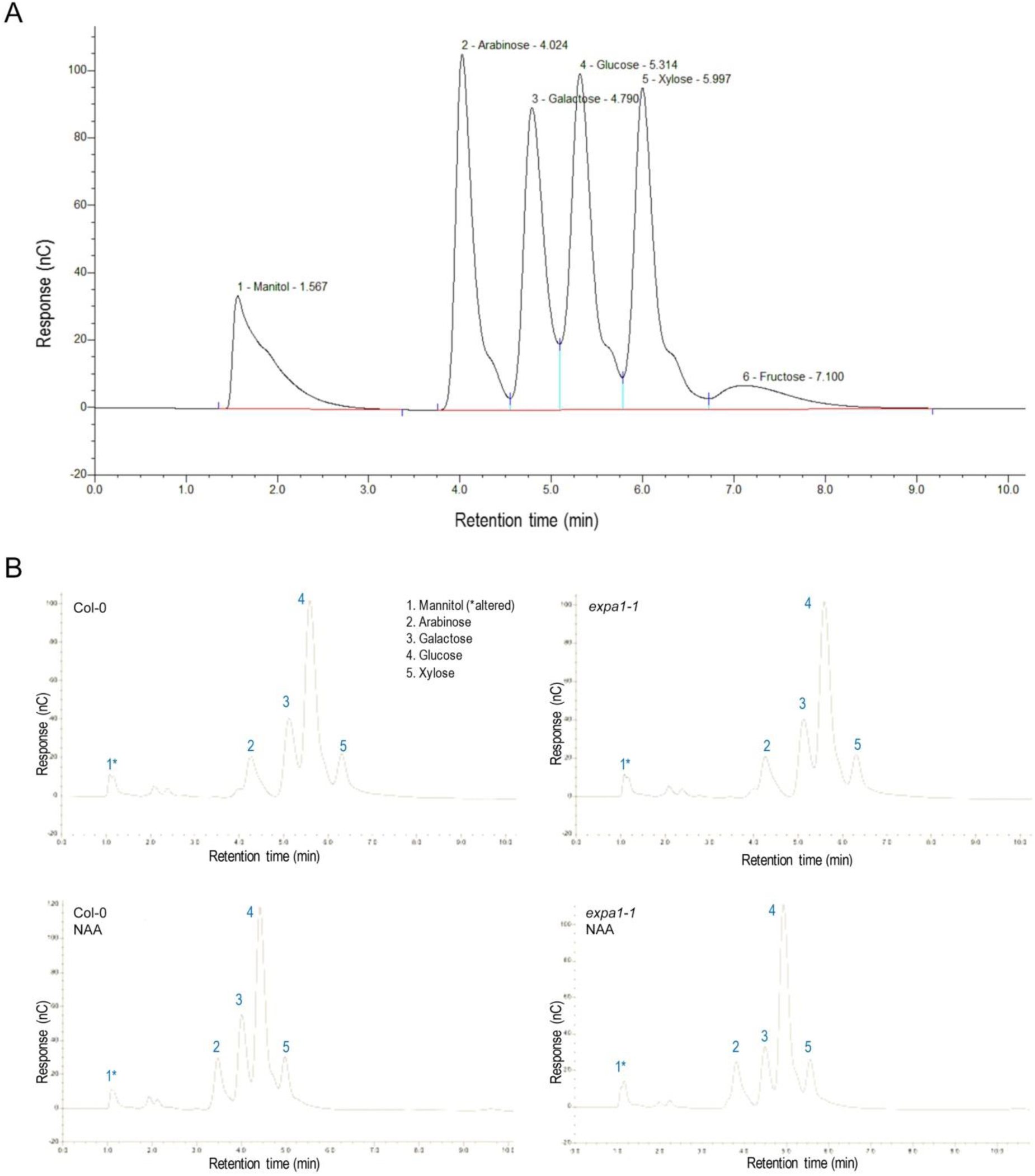
(**A**) Separation of monosaccharide sugar standards analysed by High Performance Anion Exchange Chromatography - Pulsed Amperometric Detection (HPAEC-PAD) analysis. Peaks represent the sugars present and the area under the peak allows quantification of the amount of sugar based on standards that have been run through the column first to use for quantitative comparison. **(B)** Representative chromatogram of sugar monomers of acid hydrolyzed whole roots of Col-0 and *expa1-1* with and without 6 h 10 µM NAA treatment.

**Figure S4.**
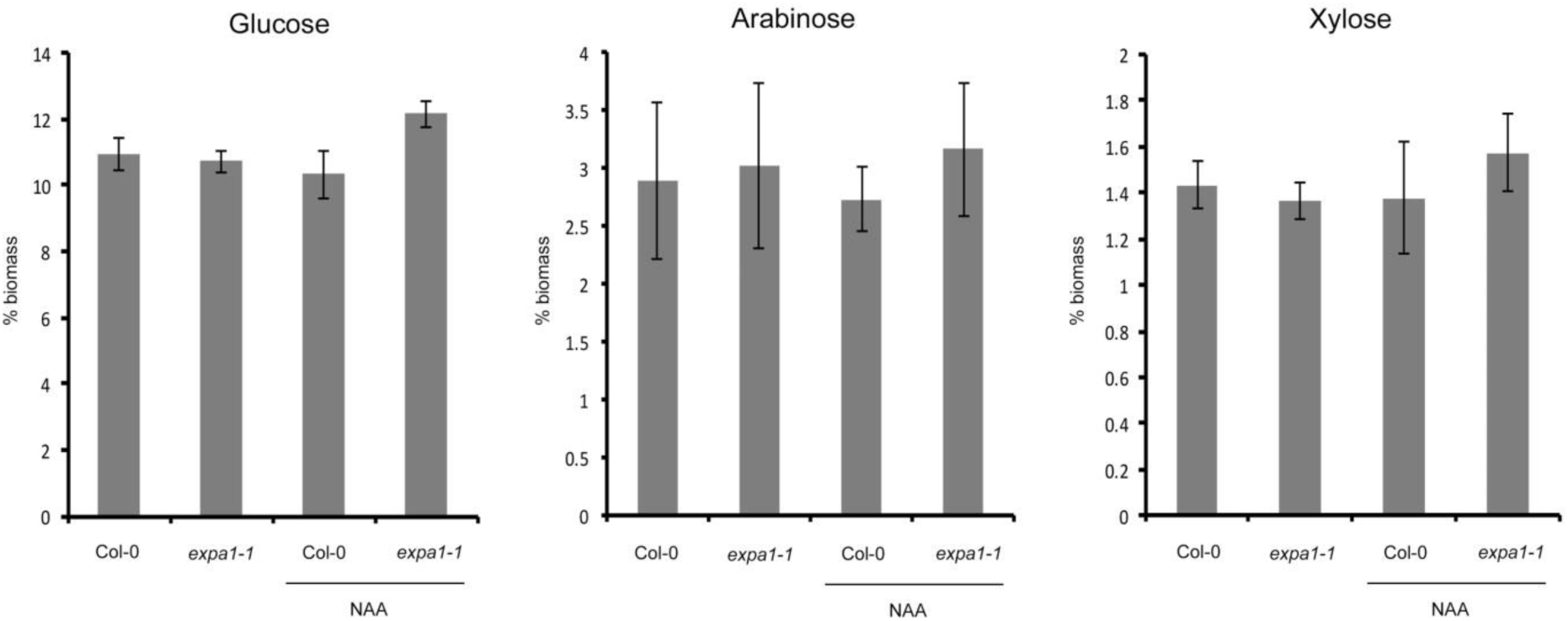
Monosaccharide sugar analysis on hydrolyzed Col-0 and *expa1-1* roots at 7 days after germination with and without 6 h of 10 µM NAA treatment using High Performance Anion Exchange Chromatography - Pulsed Amperometric Detection (HPAEC-PAD). Average of two independent biological replicates are shown, each with 2-3 technical replicates, ± standard error. There is no statistical difference (Student’s t-test) compared with Col-0 at p-value < 0.05.

**Figure S5.**
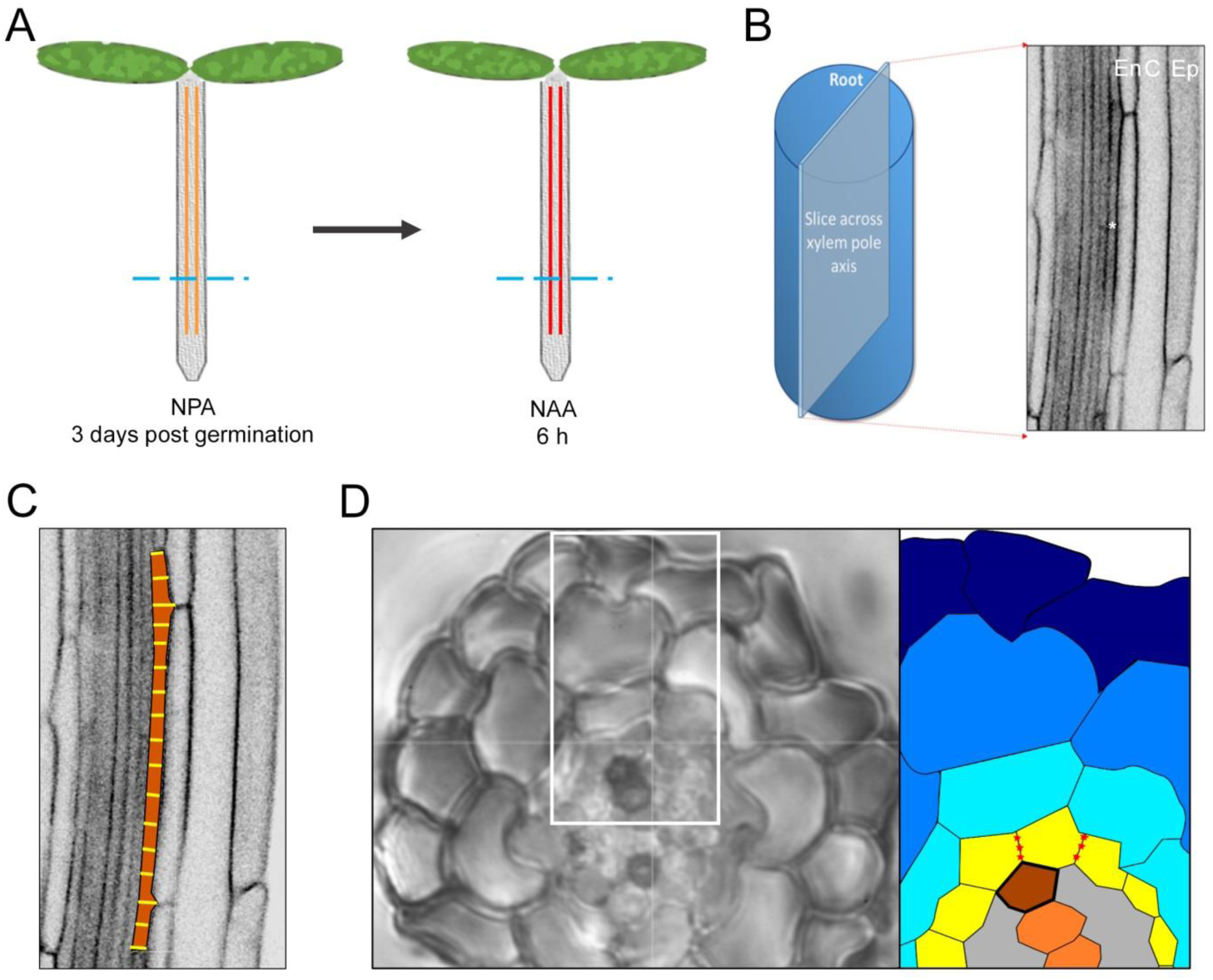
(**A**) Schematic of the lateral root induction system. Seedlings were exposed to the auxin transport inhibitor 1-N-naphthylphthalamic acid (NPA), followed by transfer of seedlings to the synthetic auxin 1-naphthaleneacetic acid (NAA) for 6 h. Vertical parallel lines represent the pericycle with red indicative of auxin induction in the pericycle that drives lateral root initiation, the dotted blue lines represent the region where the cell measurements were made (see B). **(B)** Illustration of root confocal slice used for cell width measurements. *, pericycle; En, endodermis; C, cortex; Ep, epidermis. **(C)** Indication of measurement approach to determine pericycle cell (orange) width by averaging multiple positions along this pericycle cell (yellow lines). **(D)**
*Arabidopsis* root cross section with white box marking region used for schematic. Schematic of root cross section with the pericycle in yellow and xylem pole pericycle junctions imaged marked as red stars.

**Figure S6.**
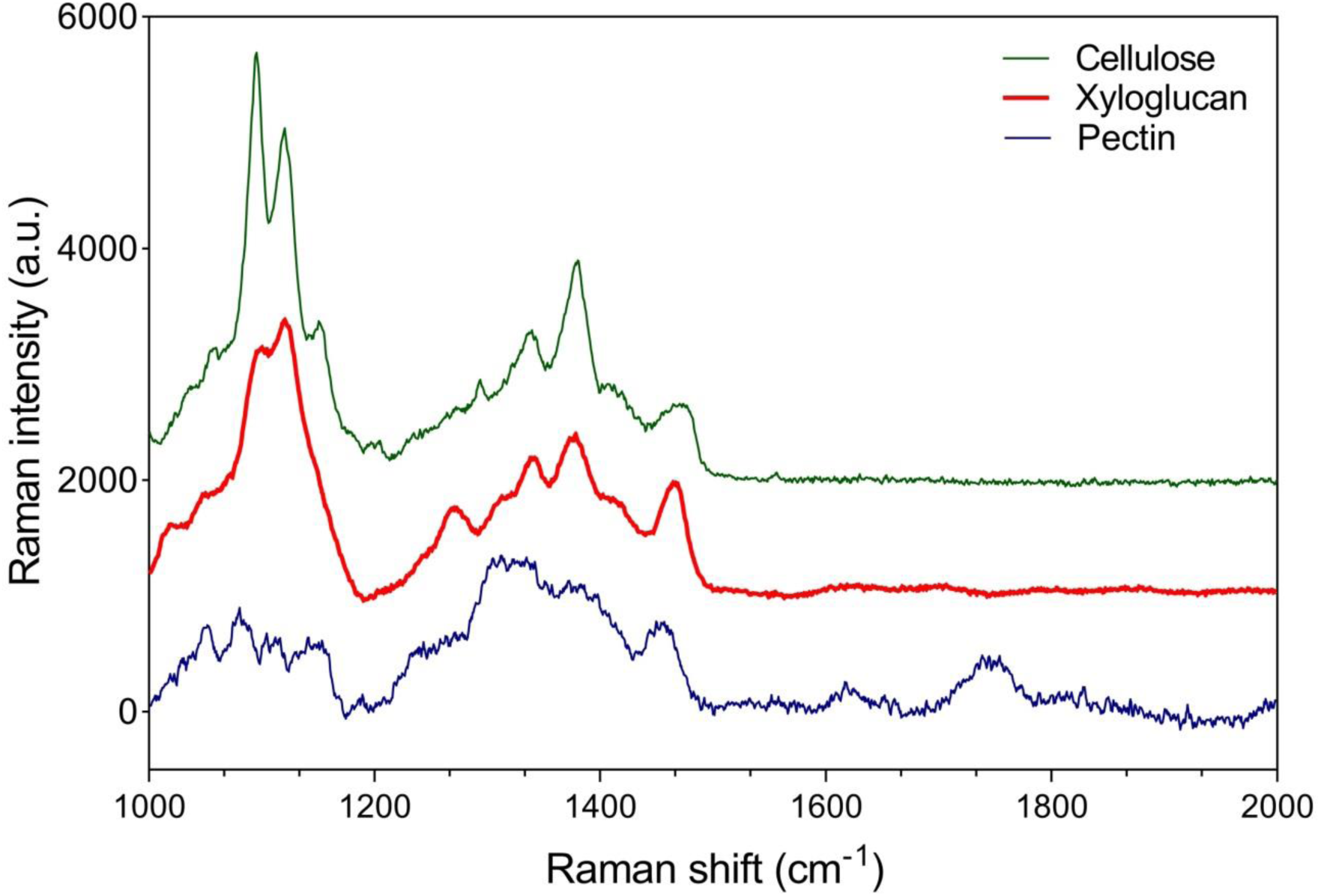
Representative Raman spectra of pure cell wall compounds used as reference spectra. The spectra of purified commercial xyloglucan and pectin were found to contain a high level of autofluorescence and so were photobleached for 30 mins prior to spectral acquisition; cellulose did not require photobleaching. a.u., arbitrary units.

**Figure S7.**
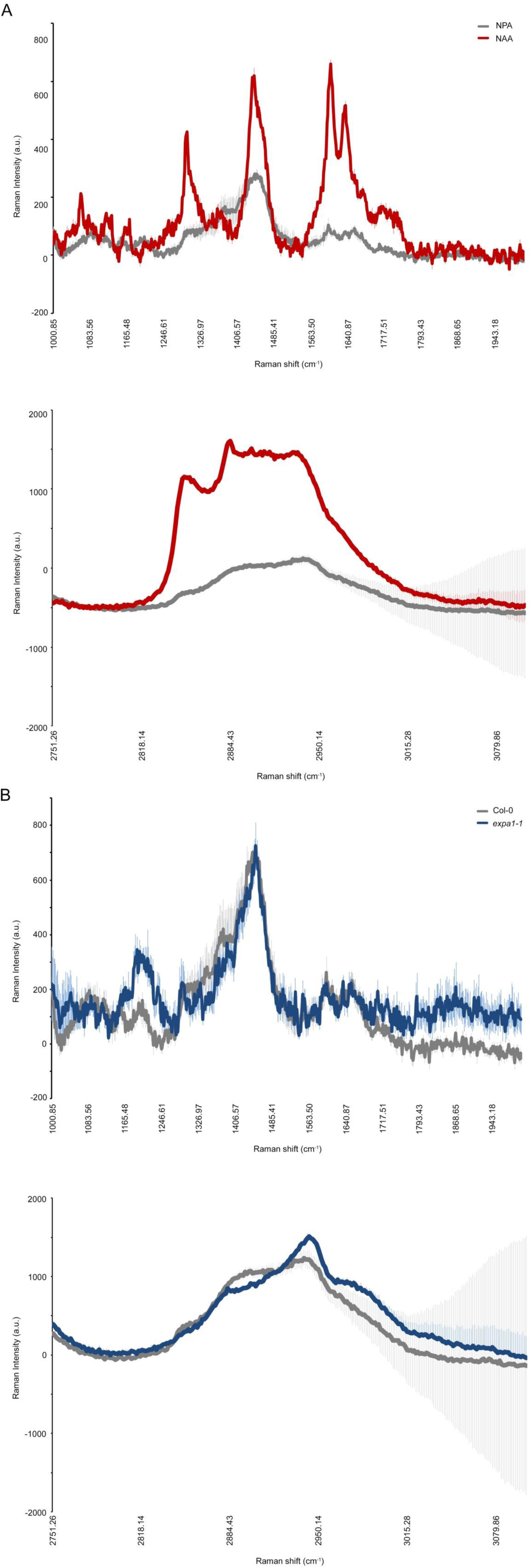
Average of 4-6 measurements ± standard error depicting chemical spectra in the range 1000 – 2000 cm^−1^ (top) and 2750 – 3100 cm^−1^ (bottom) of pericycle cell junctions on cross sections of **(A)** NPA and NAA-treated Col-0 roots and **(B)** NPA-grown Col-0 and *expa1-1* roots. a.u., arbitrary units.

**Figure S8.**
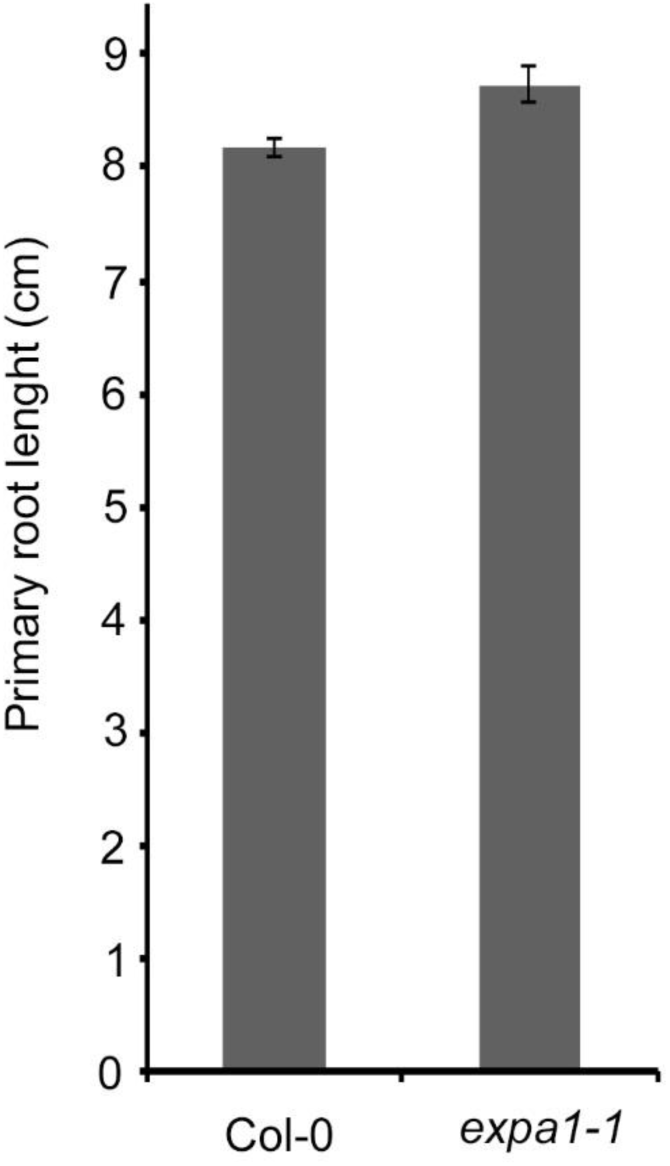
Primary root length of Col-0 and *expa1-1* at 10 days after germination. Student`st-test gave no significant difference.

**Figure S9.**
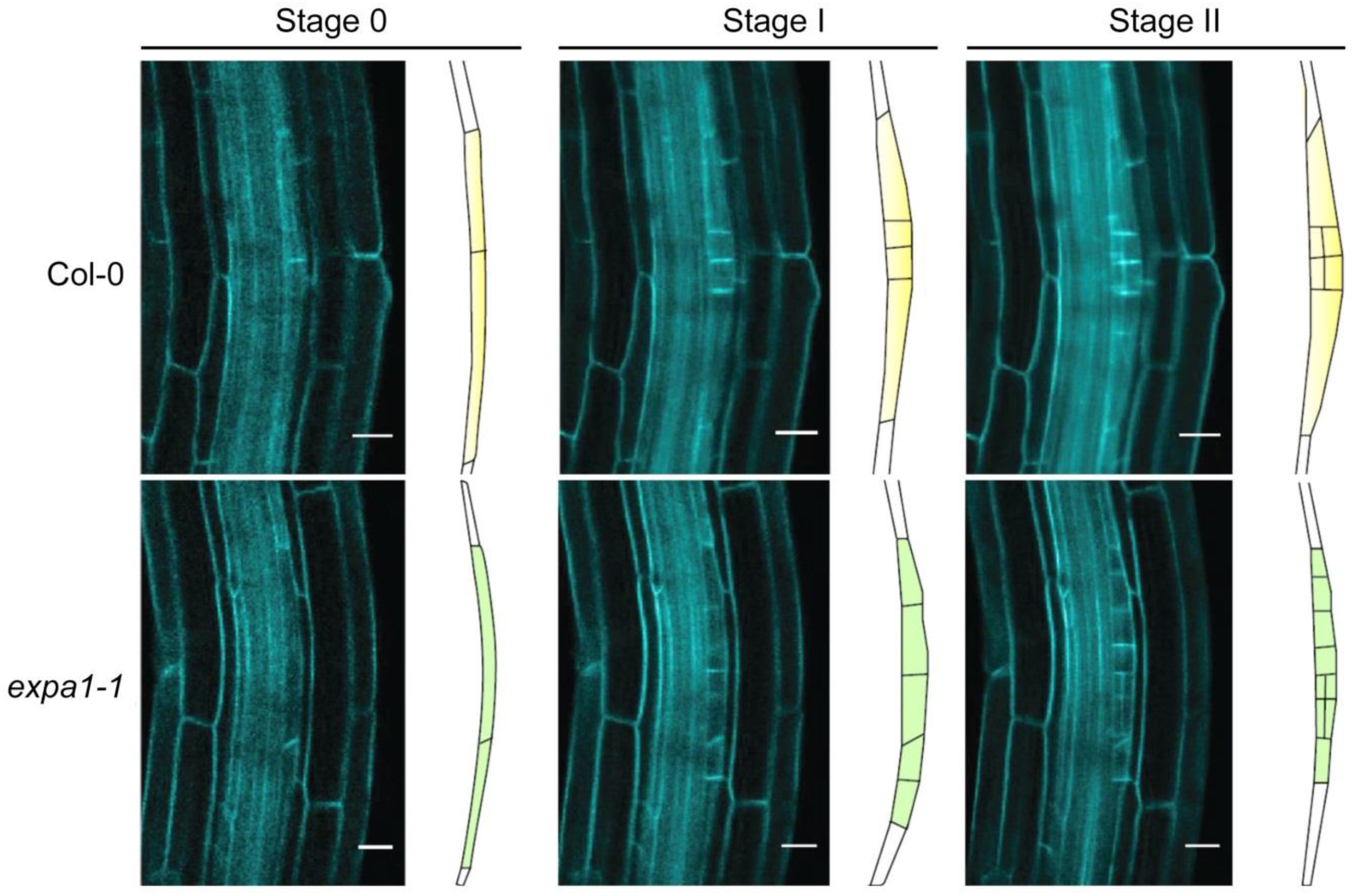
Visualization of progression through early lateral root development stages in roots of 3 days post germination seedlings that were imaged 12 h after gravitropic root bending in wild type (Col-0) and *expa1-1*. Walls were visualized using the plasma membrane marker *pUBQ10::EYFP:NPSN12* (referred to as WAVE131YFP).

**Figure S10.**
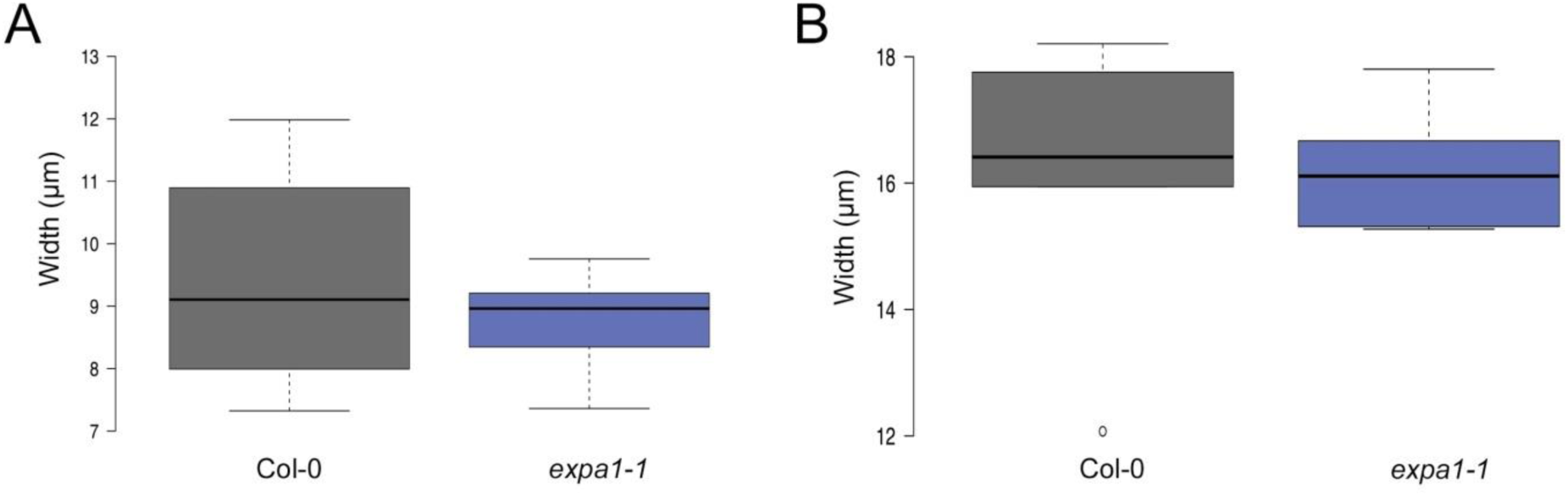
Boxplots of endodermis (A) and cortex cell width (B) in Col-0 and *expa1-1* roots of seedlings at 3 days after germination on 10 µM NPA. Data from 6-8 roots with at least 8 cells per genotype. There is no statistical difference (Student’s t-test) compared with Col-0 at p-value < 0.05.

## SI MATERIALS AND METHODS

### Plasmids and constructs

The *pEXPA1::n3xGFP* line was generated using a previously published vector backbone (58) with the *EXPA1* promoter fragment defined as 1901 base pairs upstream of the start codon. Homozygous transgenic lines were used for all experiments. For generation of expression constructs, standard molecular biology procedures and Gateway Cloning Technology (Invitrogen) was used. For the complementation construct *pEXPA1::EXPA1-6xHis*, the genomic sequence of *AtEXPA1* (fragment containing the 2.5 kb promoter, coding sequence and the UTR’s was PCR-amplified using *TTCCAAATATAGCATTGGACCGT* and *AGCACTCGAAGCACCACTT*) was cloned into the entry vector pCR8-GW-TOPO and transferred into the destination vectors pGWB7 (no promoter, 6xHis tag) (59). Plasmids were transformed into *Agrobacterium tumefaciens* strain C58 (pMP90) and *Arabidopsis thaliana* plants were transformed by floral dip as previously described (60). Homozygous, independent transgenic lines with a single insert were analyzed.

### In planta and in silico expression analysis

Total RNA was extracted from roots using a Qiagen^®^ RNeasy plant mini kit. Poly(dT) cDNA was prepared from total RNA using the SuperScript II^®^ Reverse Transcriptase (RT) from Invitrogen^®^. qPCR was performed using PerfeCTa SYBRGreen^®^ FastMix, Low ROX (Quanta) on Roche LightCycler 480 apparatus. Results of the qPCR analysis are from a minimum of two independent biological replicates with four technical replicates in each. Expression of *EXPA1* was determined using the following primers: *GATGTCAAGAAACTGGGGACA* and *GAAAGACCAGCCTGCGTTAG*. Expression levels were normalized to that of the control gene *ACTIN* using the following primers: *CTGGAGGTTTTGAGGCTGGTAT* and *CCAAGGGTGAAAGCAAGAAGA*. To assess the hormone-regulated expression of cell wall-related genes, we used Genevestigator data (on 08/07/2014) (61), retaining the experiments indicated in **Dataset S1**. The 406 putative cell wall remodeling agents were identified based on dominant families in the literature and/or information in TAIR (http://www.arabidopsis.org).

### Root and cell size measurements

The root length and number of lateral roots were determined using a dissecting microscope and ImageJ software (imagej.nih.gov/ij/). LR density indicates the number of LRs per cm of the primary root. All data are the mean values for each plant considered. Experiments were repeated three times and statistical analysis of data was performed using the Student’s *t*-test. For cell size measurement, following treatment, confocal Z-stacks of roots were collected ~400 microns from the tip with the xylem poles in focus. The stacks were processed into 2D images with the different cell layers in focus using the z-project feature in Fiji (imagej.net/Fiji). Although the curvature of the root surface varies greatly, the surface of the pericycle cells in the longitudinal confocal section is small enough to be locally flattened in 2D without too much cell shape deformation (**SI Appendix, Fig. S5**). The cell width was measured at multiple points along the cell and averaged out to account for non-uniform cell shapes of the different cell layers (**SI Appendix, Fig. S5**). To account for membrane thickness, the measurements were taken at midpoint to midpoint of the membrane surface. Boxplots were generated with shiny.chemgrid.org/boxplotr/.

### Confocal Raman microscopy

The roots were embedded in 4% Agarose (BioReagent for molecular biology, low EEO, Sigma-Aldrich) and 100-micron thick root cross sections were prepared with a Vibratome (Leica VT 1000S Vibratome). The sections were sealed in D_2_O (Sigma-Aldrich) on a glass slide with nail paint to prevent evaporation of the solvent. Confocal Raman spectroscopy and imaging was performed using a Horiba LabRAM HR microscope equipped with a piezoelectric scan stage (Märzhäuser, Germany) using either a 532 nm or a 785 nm laser, a 100x air objective (Nikon, NA = 0.9) and 50µm confocal pinhole. To simultaneously scan a range of Raman shifts, a 600 lines mm^−1^ rotatable diffraction grating along a path length of 800 mm was employed. Spectra were detected using a Synapse CCD detector (1024 pixels) thermoelectrically cooled to ~60°C. Prior to spectral acquisition, the instrument was calibrated using the zero-order line and the diagnostic phonon mode at 520.7 cm^−1^ from a Si(100) reference. Single point Raman spectra of approximately 1 and 2 micron spatial resolution (lateral and axial dimensions respectively) were acquired by optically focusing on the point of interest on the sample. The offset between the optical and laser focus was accounted for by maximizing the signal intensity as a function of sample height. For the root cross sections, Raman spectra (λ_ex_ = 532 nm) were acquired with an integration time of 120s (8 accumulations to increase the signal to noise ratio and remove spectral artefacts, e.g. cosmic rays), within the range of 1000-3200 cm^−1^, from multiple points on the pericycle cell junction and averaged to obtain the final cumulated spectra for comparative analysis (**Fig. S5D**). Root sections were taken from at least six roots of each genotype and treatment and the spectral peaks for each line and treatment were compared for consistency in spectral pattern and reproducibility of peaks. Raman spectra (λ_ex_ = 785 nm) of the reference materials were collected to use as the basis for comparative post-analysis (**Fig. S6**). All spectra were processed using LabSpec 6 software (Horiba Scientific). The spectra were baseline corrected with a local polynomial fit of the data, and subsequently smoothed using the DeNoise filter embedded in LabSpec 6 software. Spectral features within the range 2000-2750 cm^-1^ associated with the D-O stretching vibrations in D_2_O were removed from the analysis and thus only vibrational modes within the ranges 1000-2000 cm^−1^ (fingerprint region) and 2750-3100 cm^−1^ (C-H stretching region) were compared for differences in the position and intensity of spectral features.

### Monosaccharide analysis

30 mg of homogenized, dried root tissue was subject to two stage hydrolysis. 12 M sulphuric acid (1 ml) was added and the sample was incubated for 1 hr at 37°C. This was then diluted with 11 ml MilliQ^®^ water and incubated for a further 2 h at 100°C. The sugar monomer content of the supernatant was determined by High Performance Anion Exchange Chromatography with Pulsed Amperometric Detection (HPAEC-PAD). The hydrolysate was analyzed using Dionex^®^ ICS-3000 comprised of a high-pressure GD 50 gradient pump, an analytical column (Carbopac^®^ PA20, 4 mm × 250 mm). 10 mM NaOH was used as the mobile phase and the column was flushed with 200 mM NaOH between runs. All chromatographic analyses were carried out at 30°C with a flow rate of 1.0 ml/min. Samples were diluted 1:100 in 10 mM NaOH and centrifuged at 4472 g for 10 min before loading to Dionex^®^ vials. Sugar standards ranged from 0.250 to 2 g/L of arabinose, galactose, glucose, and xylose. The chromatograms for all lines and treatment have been included in the supplementary data (**Fig. S3**).

### Comprehensive microarray polymer profiling (CoMPP) analysis

Cell wall material was isolated from 7 DAG roots of wildtype and *expa1-1* after treatment with 10 µM NAA for 6 h as alcohol-insoluble residue (AIR). Frozen material was homogenized to a very fine powder in liquid nitrogen, 70% (v/v) ethanol was added to each tube and vortexed thoroughly and spun down at 14000 rpm for 10 min and the supernatant discarded. A methanol/chloroform (1/1) mixture was added to the pellet and vortexed thoroughly, spun down at 14000 rpm for 10 min and the supernatant discarded. The pellets were re-suspended in acetone and air-dried overnight. A total of about 500 roots were used for the extraction with a yield of about 10 mg AIR for each sample. These were subjected to sequential extractions as previously described (1). The extractions were performed using only diamino-cyclo-hexane-tetra-acetic acid (CDTA), and sodium hydroxide (NaOH) with 0.1% (v/v) sodium borohydride (NaBH_4_). These respectively enriched for pectin and hemicelluloses. The extracted fractions were printed as microarrays with six replicates and three dilutions and then probed with a range of primary antibodies or carbohydrate-binding modules (**Table S1)** and visualized using the appropriate alkaline phosphatase (AP) conjugated secondary antibodies before developing as described (30). The majority of monoclonal antibodies for selected cell wall epitopes was obtained from http://www.plantprobes.net/index.php and RU1 and RU2 are from INRA, Nantes, France. The arrays were scanned and mean spot signals calculated and represented as a heat map in which color intensity is proportional to numerical value.

